# Cortical 6-9 Hz Oscillation are a Reliable Biomarker of Persistent Pain in Rats

**DOI:** 10.1101/2020.01.02.893289

**Authors:** Andrew J. Furman, Charles Raver, Ying Li, Carleigh Jenne, Kathleen Hoffman, David A. Seminowicz, Asaf Keller

## Abstract

Neural biomarkers of chronic pain offer a potential tool for improving the speed of diagnosis and delivery of treatment for this devastating disease. Here, we tested the hypothesis that pain states are associated with distinct changed in cortical brain waves. We induced neuropathic orofacial pain in female rats by chronic constriction injury of the infraorbital nerve (CCI-ION). In most animals, this resulted in lasting reductions in mechanical sensitivity thresholds, and in lasting increases in facial grimace scores. We recorded electrocortigraphy (ECoG) signals over the neocortex of these rats, before and after CCI-ION, and analyzed these signals with a novel, spectral modelling approach. Consistent with our hypothesis, power in the 6-9 Hz bandwidth of the ECoG was differentially modulated in animals displaying signs of chronic pain. Specifically, development of mechanical hypersensitivity correlated with a *decrease* in 6-9 Hz power. Furthermore, we show that changes in the power of this oscillation after injury, obtained at the individual animal level, provide a more sensitive marker of pain presence than do traditional between animal comparisons of post-injury oscillatory power. Identification of animals demonstrating chronic-pain behaviors was more accurate when estimates of post-injury oscillatory power were compared against each animal’s own pre-injury baseline than when compared against post-injury power estimates from animals not developing chronic pain. These results highlight the need for establishing individual-specific, “pain-free” baselines from which oscillation disturbances can be measured and which may constitute a reliable, low-cost approach not only for diagnosing chronic pain, but also for identifying individuals likely to transition from acute to chronic pain.

---

Despite its pervasiveness, afflicting nearly one in four Americans, and resources devoted towards it, with medical costs totaling more than $600 billion annually (Gaskin & Richard, 2012), chronic pain remains exceedingly difficult to diagnose. Indeed, diagnosis can take years and multiple clinical visits, and even then, most patients fail to achieve satisfactory pain reduction (Schulte et al., 2010). One contributing factor to this crisis is that chronic pain often presents long after tissue damage has resolved, or in the complete absence of observable tissue damage. This can place physicians in an uncertain diagnostic landscape, often relying almost exclusively on patient reports, that can become even more uncertain in the face of potential malignant drug-seeking (Crowley-Matoka & True, 2012). This affects patient healthcare both directly and indirectly. Many physicians cite poor pain assessment as a major barrier to patient treatment (Von Roenn et al., 1993), while awareness of clinician distrust or inaction can negatively impact patient opinions of medical providers and dissuade future treatment seeking (Upshur et al., 2010).

Development of new diagnostic tools for chronic pain, including brain-derived biomarkers, will positively impact both patients and their medical providers. Chronic pain is associated with changes in the central nervous system that contribute to the disease’s etiology, including alterations to oscillations that are prominent in electrical recordings of the brain, like electroencephalography (EEG) (Llinás et al., 2005). These oscillations, reflecting synaptic-driven fluctuations in the membrane potentials of multiple cells (Buzsaki & Wang, 2012), are often separated into predefined bands, such as the 8-12 Hz “Alpha” rhythm (Berger, 1929) or the 4-8 Hz Theta rhythm (Kahana et al., 2001). In studies of chronic pain, attention has been focused predominantly on Alpha and Theta rhythms. Relative to age matched controls, humans with chronic pain display elevated power in both frequency bands (e.g. Sarnthein et al., 2005), and transition of the peak frequency from the Alpha to Theta range has been reported in several chronic pain conditions (de Vries et al., 2013; Kim et al., 2019). Although intriguing, the correlational nature of these findings makes it difficult to determine whether changes reflect chronic pain *per se*, whether they reflect processes that predate pain onset, or whether they have anything to do with pain at all. For example, we have recently shown that peak alpha frequency does not slow in response to a model of musculoskeletal pain that persists for multiple weeks (Furman et al., 2019). Instead, sensitivity to this model was better captured by the speed of peak alpha frequency recorded prior to model induction, suggesting that disturbances of peak frequency in chronic pain are unlikely to reflect processes directly attributable to ongoing pain.

To overcome the correlational nature of earlier human studies, preclinical rodent studies have investigated how oscillations change in a variety of injury models. Previous studies, focusing on the thalamus and somatosensory cortex, have reported increases in Theta power, without peak slowing, following injury (e.g. LeBlanc et al., 2014). These studies have focused on between-group comparisons amongst injured and sham animals. Given that confounding physiological factors, such as CSF thickness (Rice et al., 2013), can produce substantial between-subject variability in spectral power, establishing reliable ranges for “healthy” and “pathological” power characteristics may be fundamentally untenable.

One approach to overcome this limitation is to compare, *within the same individual*, estimates of EEG activity before and after injury. Here, we tested the hypothesis that oscillatory changes within an individual can be used as a biomarker of chronic pain.

## Materials and Methods

### Animals

All procedures were conducted according to Animal Welfare Act regulations and Public Health Service guidelines. Strict aseptic surgical procedures were used, according to the guidelines of the International Association for the Study of Pain and approved by the University of Maryland School of Medicine Animal Care and Use Committee. A total of 13 female (aged 8 – 15 weeks) Sprague Dawley rats weighing 180 – 300g were used. Animals were housed in polycarbonate cages at room temperature (23 ± 0.5 °C) on a 12 h light/dark cycle (lights on from 7:00 am to 7:00 pm), and allowed access to standard chow and drinking water *ad-libitum* throughout the study. To avoid potential damage to implants, rats were single housed.

### Electrocorticographic (ECoG) Recordings

We anesthetized animals continuously during surgery with inhalant isoflorane, and placed them in a stereotaxic device, while monitoring body temperature with a thermoregulated heating pad. We implanted a differential, telemetric transmitter (ETA-F10, Data Science International, St. Paul, MN) subcutaneously along the animal’s flank, and attached the transmitter’s leads, through 1 mm craniotomies, over the dura at coordinates for the anterior cingulate cortex (0.5 mm left of midline, 1.75 mm anterior to Bregma), and at a reference position over the cerebellum (1.5 mm lateral to midline, 4 mm posterior to Lambda).

### Chronic Constriction Injury of the Infra-orbital Nerve (CCI-ION)

As in our previous studies, we used a rodent model of neuropathic pain, evoked by chronic constriction of the infraorbital nerve (CCI-ION; Benoist et al., 1999; Akintola et al., 2017). We anesthetized animals with ketamine and xylazine and performed intra-oral surgery under aseptic conditions. Briefly, we made an incision along the gingivobuccal margin, beginning distal to the first molar, freed the ION from surrounding connective tissue, and loosely tied the nerve using silk thread (4–0), 1–2 mm from the nerve’s exit at the infraorbital foramen. All lesions were performed to the right ION to ensure recordings were taken from the ACC contralateral to injury.

### Mechanical sensitivity

A series of calibrated von Frey filaments was applied to the orofacial skin, at the cutaneous site innervated by the ION. An active withdrawal of the head from the probing filament was defined as a response. We used the up-down method to determine withdrawal thresholds, as described previously (Chaplan et al., 1994; Akintola et al., 2017). If an animal withdrew from the smallest (0.6g) or heaviest (15g) probe on three consecutive trials, the weight of the applied probe was recorded as the withdrawal threshold.

For two animals, baseline von Frey data was not available. For these animals, baseline data was defined as the average, baseline mechanical threshold of all other animals for that side of the face.

### Facial grimace

As in our previous studies (e.g., Akintola et al., 2017) we scored facial grimace by video recording animals in a Plexiglas chamber, for a period of 20 min. Scoring the facial expressions is a semi-automated procedure that uses the “Face Finder” application (Sotocinal et al., 2011)— provided to us by J.S. Mogil—to select images for scoring. The grimace scale quantifies changes in four “action units”, orbital tightening, nose-cheek bulge, whisker tightening and ear position for rats. We screened, labeled, randomly scrambled and scored these images, with the experimenter blinded to the treatment groups and identity of each image. We selected eight to ten images for each animal at each time point. For each image, each of the four action units was given a score of 0, 1, or 2, as previously described (Akintola et al., 2017; Sotocinal et al., 2011). We computed mean grimace scores as the average score across all the action units.

### Animal activity

Animal activity during ECoG recording was captured by continuously (at 250Hz) measuring the strength of the transmitter signal. Because the strength of the transmitter signal is determined by its distance from a fixed point on the receiver, movement of the animal in the vertical or horizontal dimension result in either increases (moving towards the receiver) or decreases (moving away from the receiver) in signal strength; for an animal that is immobile, transmitter signal strength will remain stationary. Animal activity can thus be approximated by summing the absolute, first derivative of the transmitter signal. For the purpose of analysis, signal strength data were segmented into 15 second epochs. Only epochs in which ECoG data was analyzed were included in relevant analyses.

### Procedures

An overview of the experimental procedures is provided in Figure 1. Starting one week prior to CCI-ION, and one week after ECoG surgery, we handled and acclimated the animals to all experimental apparatuses, on three alternating days, to reduce anxiety and stress. We collected baseline facial grimace and mechanical sensitivity scores on the morning before to CCI-ION surgery. We collected facial grimace data also after CCI-ION, on post-surgery days 7, 14 and, 21; and collected mechanical sensitivity scores on post-surgery days 7 and 21.

**Figure 1.**
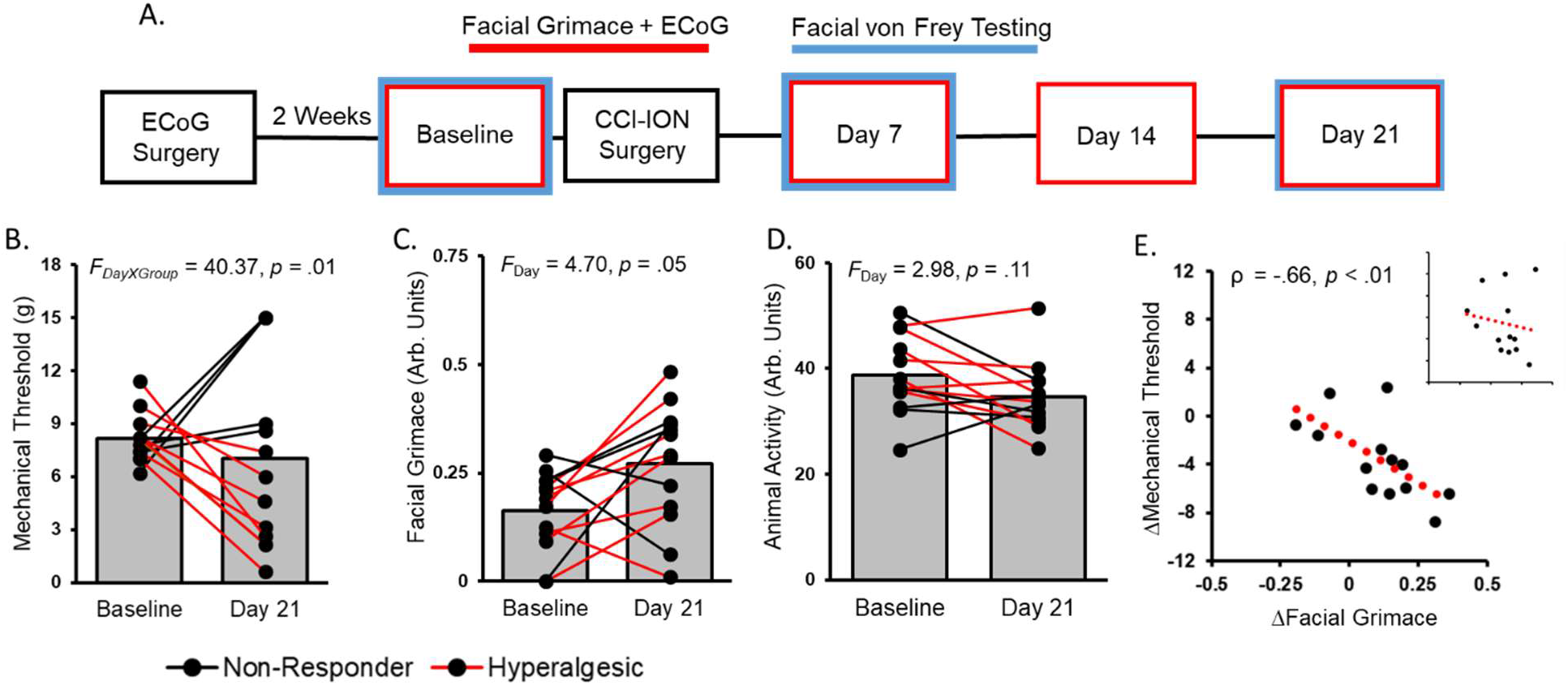
Twenty-one days after CCI-ION, mechanical sensitivity on the right side of the face, and facial grimace scores signal the presence of chronic pain. **A**. Timeline of experimental procedures. Measurements from the Baseline timepoint were collected on the same day as CCI-ION surgery was performed. Red and blue bounding boxes reflect timepoints where facial grimace or mechanical sensitivity were tested, respectively. **B**. Mechanical sensitivity recorded from the right side of the face varies based on group type. Black lines reflect data from non-responder animals and red lines reflect data from hyperalgesic animals. **C**. Facial grimace increases following CCI-ION, regardless of group membership. **D**. Animal activity does change following CCI-ION regardless of group membership. **E**. Changes in mechanical sensitivity and facial grimace (Day 21 – Baseline) are correlated when accounting for changes to mechanical sensitivity on the left side of the face in non-responder animals. Shown in the inset is the correlation when only sensitivity from the right side of the face is used. Red lines reflect the linear regression line of best fit.

For a subset (n=5) of animals, this protocol was first completed before CCI-ION and then again after CCI-ION. Within cohorts of animals, naïve and injured animals were tested simultaneously, and animal identity was unknown to experimenters until completion of all procedures.

## Analysis

### ECoG Processing

The primary data of interest was ECoG collected during the first fifteen minutes of Baseline and during Day 21 facial grimace testing. Inspection of facial grimace recordings revealed that this time period was largely absent of periods of sustained immobility or sleep across animals.

We processed ECoG data using the Chronux toolbox (version 2.11, http://chronux.org/) in MATLAB (Mathworks, Natick, MA). We visually inspected raw data to ensure quality of recording and then segmented the data into 15-second epochs. We discarded epochs with maximum amplitudes greater than 3 standard deviations above the mean amplitude across all epochs for a given animal. On average, out of the total 60 epochs per time point, 7.8 (range: 2 – 14) and 7.0 (range: 3- 12) epochs were discarded per animal from Baseline and Day 21 recordings, respectively. We converted accepted epochs to the spectral domain using a multi-taper approach, with 10 tapers and a time-bandwidth product of 5. Finally, spectral data was log_10_ transformed and then Z-scored to normalize data across animals and sessions.

### Spectral Quantification

Traditional spectral analysis of human or primate electrophysiological data often relies on segregating spectral power into pre-defined frequency bands such as Theta (4-8 Hz) or Alpha (8-12 Hz). While useful, this approach is limited in its application to the current study, given that the boundaries of these canonical frequency bands are not well established in rodents. Indeed, inspection of spectra revealed that power was predominantly expressed across a frequency range encompassing both the Theta and Alpha bands (see Figure 2B). To best capture this range of activity, we elected to empirically define the frequency range of this oscillation using a spectral modelling approach similar to that used in MR spectroscopy (Vanhamme et al., 1997). In brief, this entails removal of the 1/f “broadband” or “aperiodic” activity from the spectra, and then modelling appropriately sized residuals with a Gaussian probability density function. This approach not only provides an opportunity to objectively define a spectral range of interest, but also ensures that the influence of aperiodic activity, which can distort power estimation, is minimized (Haller et al., 2018).

**Figure 2.**
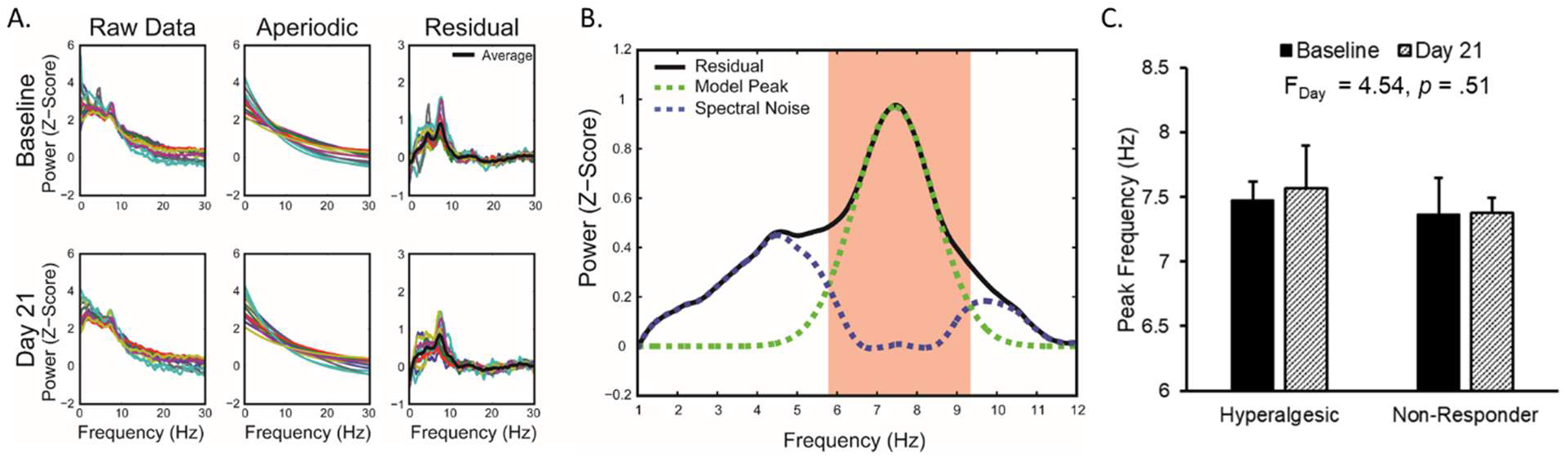
Spectral power is primarily expressed in a 6-9 Hz oscillation. **A.** Overview of aperiodic modelling. For each animal, the average spectra on each day was normalized with log10 transformation followed by z-scoring (left panel). The aperiodic, shown in the center panel, was then fit using an exponential function. The fit aperiodic was then subtracted from the raw data to yield a spectral residual (right panel). Each colored line represents a single animal and color-animal pairing is preserved across panels. **B.** The oscillation of interest was identified by fitting a Gaussian probability density function to the peak appearing at ~7 Hz in the average Baseline residual (green line). The fit peak was then subtracted from the original residual to yield a second “noise” residual (blue-dotted line). The oscillations range was defined as all frequency bins where the estimated power of the modelled peak surpassed the noise residual (shown as a shaded red zone). **C.** The estimated peak frequency within this range does not show slowing after CCI-ION.

For each animal’s average Baseline and Day 21 spectra, we used data in 0.1-1 Hz and 15-30 Hz ranges—frequency spans where no narrowband spectral activity was present—to perform a least-squared regression fit of the aperiodic with an expanded exponential equation, including variables for the amplitude (*a*), exponent (*b*), and vertical shift (*c*):

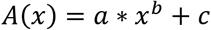

We then subtracted the modeled aperiodic from the original data to yield an “aperiodic-free” set of spectral residuals. We plotted all aperiodic fits against the original data to ensure proper fitting.

For Baseline spectra only, we averaged residuals across animals to yield a grand average residual that was then used to identify spectral peaks, or “narrowband” activity in the data. We elected to focus only on Baseline spectra in order to identify a spectral range of interest that bears no *a priori* relationship to pain. This, in our estimation, provides the most unbiased approach for determining the impact of pain on spectral activity.

We then used the standard deviation of the grand average residual to determine an empirical threshold, 3 standard deviations above the mean of the residual, for identifying spectral peaks. Least squares regression was then used to fit identified peaks to a Gaussian probability density function, G*(x)*:

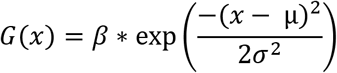

Where β determines the height of the Gaussian, μ determines the position of its center, and σ determines its width. To avoid fitting multiple peaks with a single Gaussian, we defined the fitting range of an identified peak as the range of consecutive frequency elements (i.e. 8 Hz, 8.1 Hz, 8.2 Hz,…) where the slope uniformly increases up to the peak and then uniformly decreases after passing the peak.

We subtracted modeled Gaussians from the original grand average residuals to produce a second residual representing spectral noise. We then identified our range of interest for a given peak as the frequency range where the contribution of the modelled Gaussian to the first residual is greater than that contributed by the estimate of spectral noise (see shaded area of Figure 2B).

### Statistical Analysis

All analyses were performed using custom scripts implemented in the Matlab environment (version R2013A). Statistical tests were conducted in Matlab or SPSS (Version 25).

Animals were classified as “hyperalgesic” if mechanical sensitivity testing revealed a minimum 15% decrease in mechanical threshold. Animals not meeting this criterion were classified as “non-responders”.

We first investigated whether changes in mechanical sensitivity (for the right side of the face; ipsilateral to CCI-ION) occurred after CCI-ION by comparing scores on Baseline and Day 21 testing sessions with a 2X2 factorial ANOVA with Day (Baseline vs. Day 21) and Group (Hyperalgesic vs. Non-Responder) as within and between-subject variables, respectively. All *post-hoc* contrasts were Bonferroni corrected, yielding a significance threshold of .0125. An identical ANOVA was then performed for both facial grimace and animal activity data. The relationships of mechanical sensitivity, grimace scores, and animal activity were investigated with Spearman rank-order correlations.

To determine whether power changes in a spectral range of interest occur following CCI-ION, we submitted the mean power estimate for each animal to a 2X2 factorial ANOVA with Day (Baseline vs. Day 21) and Group (Hyperalgesic vs. Non-Responder) as within and between-subject variables, respectively. All *post-hoc* contrasts were Bonferroni corrected, yielding a significance threshold of .0125.

To determine if changes in spectral power were related to behavioral metrics, we correlated changes in animal activity, mechanical sensitivity, and grimace scores with spectral power changes by using separate Spearman rank-order correlations. To account for the possibility that differences in changes to mechanical thresholds on the two sides of the face, ΔMechanical_R-L_, impact the relationship of facial grimace changes, ΔFacial Grimace, to spectral power changes, ΔPower, we regressed ΔMechanical_R-L_ and ΔFacial Grimace and against ΔPower.

We tested the significance of spectral changes occurring within individual animals. For each animal, and independently for Baseline and Day 21 data, we subtracted the aperiodic fit obtained from that animal’s grand mean spectra from the spectra of each individual epoch. This yielded a set of residuals for each animal on each day, from which epoch-level spectral power could be calculated. We compared distributions of spectral power from Baseline and Day 21 by calculating area under the curve (AUC) using nonparametric receiver operating characteristics (Fawcett, 2006). To assess significance, we employed a bootstrapping approach where labels (Baseline or Day 21) were randomly assigned to spectral estimates to create a null dataset from which AUC could be calculated. This process was repeated 10,000 times to generate a distribution of null AUC scores. Obtained AUC values were deemed significant if they surpassed the 95th percentile of the null distribution. We assessed the pattern of identified changes across groups using Fisher’s exact test on a 2 (Hyperalgesic vs. Non-Responder) X 3 (power increase, no change, power decrease) contingency table.

Finally, we compared whether changes in spectral power or actual power amounts at Day 21 were better indicators of chronic pain. Power changes were calculated by subtracting the average 6-9 Hz power at Baseline from each Day 21 epoch estimate. For each metric, values were pooled across animals and labelled as having come from a hyperalgesic or non-responder animal. Traditional ROC statistics were then calculated for each metric and significance was assessed by determining whether obtained AUC values surpassed the 95^th^ percentile of a bootstrapped, null distribution. Finally, the significance of the difference between obtained AUC values for power change and power amount was assessed against a bootstrapped distribution of null, absolute AUC differences.

## Results

Twenty-one days after CCI-ION, changes in mechanical sensitivity thresholds on the injured, right side of the face were used to define two groups of animals: 1) Animals experiencing threshold decreases relative to their Baseline values (n = 8; Mean_Baseline_ = 8.66g, 95% CI = [7.67, 9.65]; Mean_Day21_ = 3.59g, 95% CI = [2.03, 5.15]) and 2) Animals demonstrating no change, or increases in threshold (n = 5; Mean_Baseline_ = 7.31g, 95% CI = [6.65, 7.97]; Mean_Day21_ = 12.52g, 95% CI = [9.54, 15.5]). Increases in mechanical thresholds after CCI likely represent nerve damage resulting from inadvertent, excessive constriction of the nerve (Vos et al., 1994; Akintola et al., 2017). We refer to the former group of animals as “hyperalgesic” and to the latter as “non-responders”. A 2X2 factorial ANOVA with Day (Baseline vs. Day 21) and Group (Hyperalgesic vs. Non-Responder) as within and between-subject variables, respectively, revealed an effect of Group, F_(1,11)_ = 17.98, *p* < .01, partial η^2^ = .62, that was qualified by the Day X Group interaction, F_(1,11)_ = 40.37, *p* < .01, partial η^2^ = .79. No main effect of Day was found, F_(1,11)_ = .01, *p* = .94. Not surprisingly, the Day X Group interaction revealed that thresholds increased for non-responder animals and decreased for hyperalgesic animals (both *p* < .01). Interestingly, mechanical sensitivity thresholds on the left side of the face decreased equally for both animal groups (n = 13; Mean_Baseline_ = 8.43g, CI = [7.67, 9.19]; Mean_Day21_ = 5.39g, CI = [3.90, 6.88], Day: F_(1,11)_ = .12.79, *p* < .01, Group: F_(1,11)_ = .82, *p* = .39,). Note that the Day X Group interaction showed that hyperalgesic animals (Mean = −4.22) had larger changes than non-responder animals (Mean = −1.12), F_(1,11)_ = 4.29, *p* = .06.

During this same time period, facial grimace scores also increased (n = 13; Mean_Baseline_ = .16, 95% CI = [.11, .21]; Mean_Day21_ = .27, 95% CI = [.20, .34], Figure 1C). A 2X2 factorial ANOVA revealed an effect of Day, F_(1,11)_ = 4.70, *p* = .05, but not Group, F_(1,11)_ = .40, *p* = .54, or Day X Group, F_(1,11)_ = .41, *p* = .54, indicating that both groups of animals underwent similar changes in facial grimace. Changes in facial grimace scores were not correlated with changes in mechanical sensitivity thresholds from the right side of the face, Spearman ρ = −.22, *p* = .46 (Figure 1D inset). One possible explanation is that facial grimace is a composite measure that captures pain reflected on either side of the face; for non-responder animals, the above correlation may fail to capture threshold changes occurring on the left side of the face. Indeed, using mechanical sensitivity thresholds from the left side for non-responders—the side that shows hyperalgesia—revealed a clear relationship between changes to facial grimace and changes to mechanical sensitivity, Spearman ρ = −.66, *p* = .01 (Figure 1D).

Importantly, animal activity—assessed from signal strength recorded from the implanted transmitters (see Methods)—did not appear to change between the two recording sessions (n = 13; Mean_Baseline_ = 34.08, 95% CI = [33.64, 42.53]; Mean_Day21_ = 34.44, 95% CI = [28.70, 40.20], Figure 1D). Neither of the main effects, Day: F_(1,11)_ = 2.98, *p* = .11; Group: F_(1,11)_ = 1.18, *p* = .30, nor the Day X Group interaction, F_(1,11)_ = .87, *p* = .37, reached significance.

### A 6 – 9 Hz Oscillation Decreases Following CCI-ION

ECoG spectra recorded during Baseline facial grimace testing revealed putative narrowband activity occurring across a frequency range encompassing both Theta and Alpha frequencies (Figure 2). Spectral modelling identified a single spectral peak, μ, at 7.44 Hz (α = .97, σ = 1.00) that was estimated to contribute most of the power to spectral residuals in a range of interest (ROI) extending from 5.78 Hz to 9.34 Hz (Figure 2B). To avoid missing oscillations that may disappear or be reduced after injury, only data from Baseline spectra were used to identify this ROI.

Analysis of ROI spectral power revealed a significant Day X Group interaction, *F*_(1,11)_ = 11.72, *p* < .01, partial η^2^ = .52 (Figure 3A). Whereas non-responder animals experienced power gains in our ROI from Baseline to Day 21 (n = 5; Mean_Baseline_ = .59, CI = [.40, .78]; Mean_Day21_ = .70, CI = [.52, .88]), hyperalgesic animals experienced power decreases (n = 8; Mean_Baseline_ = .73, CI = [.53, .93]; Mean_Day21_ = .55, CI = [.34, .76]). Neither the main effect of Day, *F*_(1,11)_ < .01, *p* = .97, nor the main effect of Group, *F*_(1,11)_ = .59, *p* = .46, reached significance. Simple effect contrasts showed that the change in power was significant for hyperalgesic (*p* < .01) but not non-responder animals (*p* = .12). There were no differences between groups at either Baseline (*p* = .36) or Day 21 (*p* = .33).Importantly, changes in animal activity—that may confound interpretation of changes in ECoG signals—were not correlated with oscillatory changes, Spearman ρ = .22, *p* = .47 (Figure 3B).

**Figure 3.**
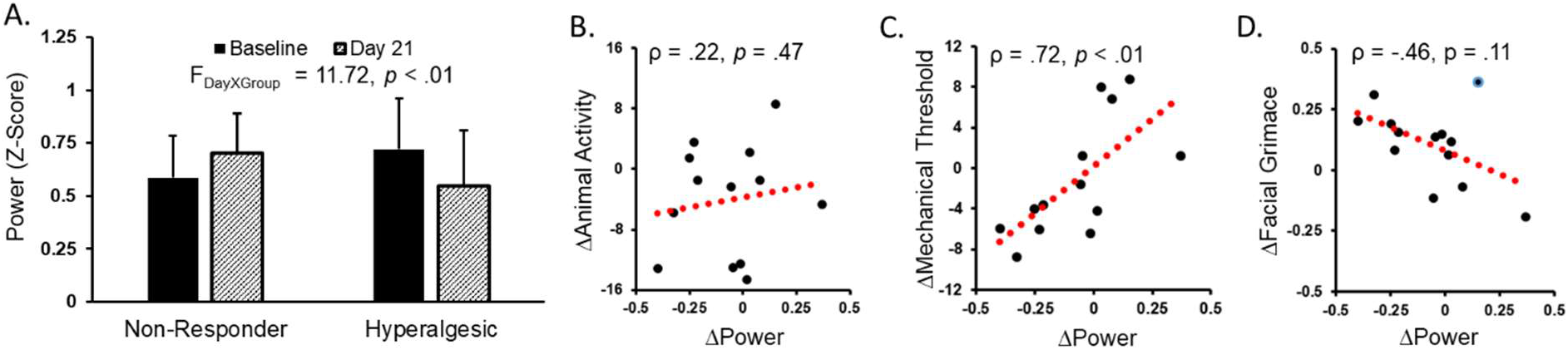
Spectral power in the identified 6-9 Hz oscillation is modulated based on the behavioral outcome of CCI-ION. **A.** Mean estimates (+1 S.D.) of 6-9 Hz power for each group on each day. Whereas power decreases for hyperalgesic animals, power increases for non-responder animals. **B.** Changes in animal activity are not related to changes in spectral power. **C.** Changes in power are positively correlated with changes in mechanical sensitivity. **D.** Changes in power are only moderately related to changes in facial grimace. This relationship appears to be weakened by the presence of an outlier, highlighted in blue. Removal of this animal increases the magnitude of the relationship (Spearman ρ = −.77). Dotted red lines reflect the linear regression line of best fit.

We found no evidence of oscillation *slowing* following CCI-ION, with nearly identical peak frequencies recorded at Baseline (n = 13; mean = 7.43, CI = [7.32, 7.54]) and Day 21 (n = 13; mean = 7.49, CI = [7.34, 7.64]; Figure 2C). Analyzing the data with a 2 × 2 factorial ANOVA including Day (Baseline vs. Day 21) and Group (Hyperalgesic vs. Non-Responder) as within and between subject variables, respectively, revealed that neither main effect nor the interaction reaches significance (all *F* < 1.77, all *p* > .21).

Significantly, changes in ROI spectral power were strongly correlated with changes in mechanical thresholds on the right side of the face, Spearman ρ = .72, *p* < .01, but not the left side, Spearman ρ = .35, *p* = .24 (Figure 3C). Changes in facial grimace were only moderately related to power changes, Spearman ρ = −.46, *p* = .11, but this lack of relationship appeared to be due the presence of one data point with excessive leverage (ρ = −.77 without data point; Figure 3D). Notably, this non-responder animal had experienced a large decrease in threshold on the left side of its face. Given that grimace captures pain responses occurring on either side of the face, we accounted for differences in threshold change between the two sides of the face by regressing power changes against changes to facial grimace scores and against the difference between mechanical threshold changes on each side of the face (ΔMechanical_Right_ − ΔMechanical_Left_; ΔMechanical_R-L_); for ΔMechanical_R-L_, a score of 0 indicates similar changes across both sides of the face, while increasingly larger positive and negative values indicate relatively greater threshold increases or decreases on the right side of the face, respectively. The overall regression model was significant, F_(2,10)_ = 6.70, *p* = .01, R = .76, Adj. R^2^ = .49, as were the terms for ΔFacial Grimace, *t* = −2.73, *p* = .02, β = −.57, and Δ_R-L_Mechanical, *t* = −2.69, *p* = .02, β = −.56. Thus, changes in facial grimace are significantly related to changes in 6-9 Hz power after CCI-ION. Expanding the model to include a term for changes in animal activity did not change this finding for either ΔFacial Grimace, *t* = − 2.65, *p* = .03, β = −.55, and Δ_R-L_Mechanical, *t* = −2.72, *p* = .02, β = −.76, and again demonstrated that changes in animal activity, ΔAnimal Activity, are unrelated to spectral power changes, *t* = −1.07, *p* = .31, β = −.301.

### 6-9 Hz Power Decrease is a Reliable Marker of Hyperalgesia

We compared the distributions of epoch-level ROI power, to test the prediction that these are correlated with behavioral hyperalgesia. On average, Day 21 and Baseline power differed by .12 (S.D = .16) and −.18 (S.D. = .15) for non-responder and hyperalgesic animals, respectively; AUC averaged .64 (S.D = .14) and .69 (S.D. = .15) for non-responder and hyperalgesic animals, respectively. From the non-responder group, three animals (60%) demonstrated increases in ROI power, whereas none demonstrated decreases in ROI power. Conversely, five hyperalgesic animals (62.5%) had significant decreases in ROI power, with no animals showing increases. Data from a representative hyperalgesic animal is shown in Figures 4 and 5. For this animal, the obtained difference between Day 21 and Baseline, −.32, yielded a significant AUC value, .86 (95th% = .62).

**Figure 4.**
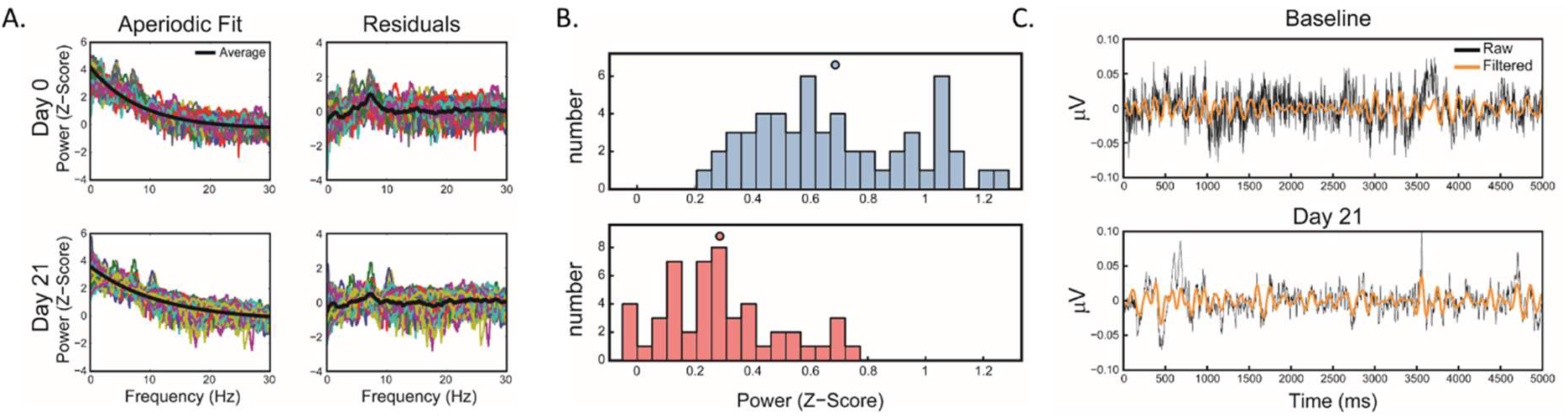
Application of spectral modelling to individual epochs. Data from one animal is shown in this example. **A.** Epochs from each day were log_10_ transformed and z-scored (left panel). The aperiodic calculated from average spectra (dotted black line) was then subtracted from each epoch to yield a set of residuals (right panel). Power was calculated for each epoch in the frequency range of the identified oscillation. Each colored line reflects a single epoch. The average across all residuals is shown by the solid black line. **B.** The distribution of 6-9 Hz power estimates for this animal on each day. Circles above each histogram reflect the mean. Less power is consistently evident on Day 21. **C.** Example raw (black) and 6-9 Hz filtered (orange) ECoG traces from Baseline and Day 21 recordings. Provided data reflect the first five seconds from epochs corresponding to the median 6-9 Hz power on their respective day. The periodicity of oscillations in the 6-9 Hz filtered signal is noticeably reduced (i.e. number of complete cycles) on Day 21.

**Figure 5.**
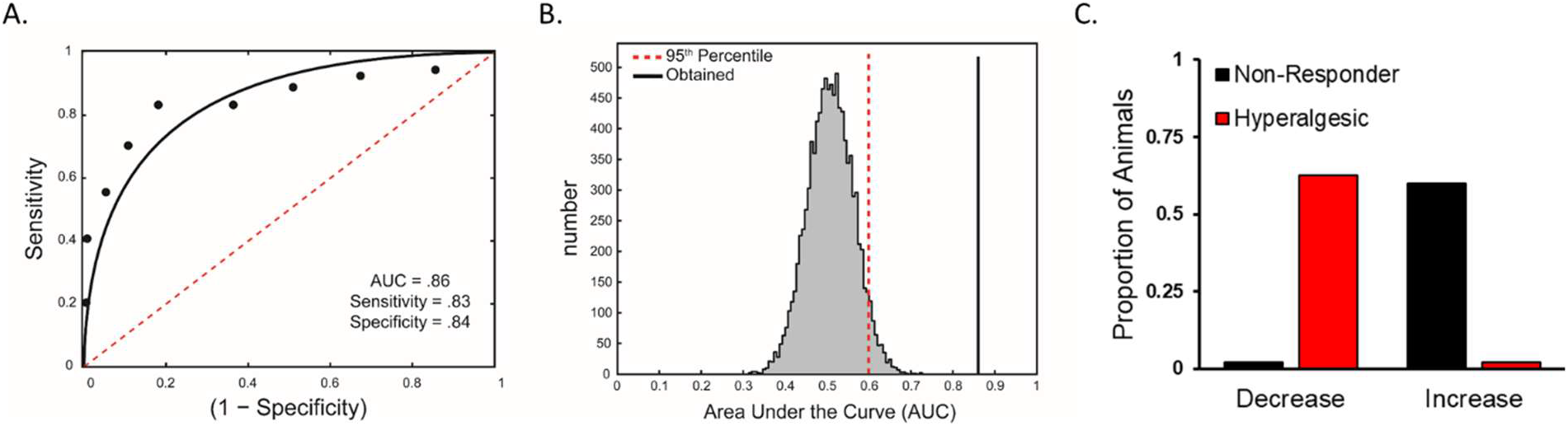
Example of null distribution testing used to determine significant within-animal changes after CCI-ION. Data shown in A and B are from the same animal used in Figure 4. **A.** Baseline and Day 21 spectral power can be reliably distinguished for this animal and (**B**) at above chance levels according to a null distribution of AUC values. **C.** All animals demonstrating significant decreases in power were hyperalgesic while all animals with significant increases in power were non-responders.

Across all animals, the average threshold for detecting a significant change in spectral power was +/− .12 (S.D. = .03). According to a Fisher’s exact test, this pattern of increases, decreases, or no change in ROI power across the two groups was significantly different from what would be expected by chance, *p* = .02. These findings suggest that a decrease in 6-9 Hz power may be a reliable marker of pain.

### 6-9 Hz Power Changes Are a Better Indicator of Chronic Pain than Day 21 Power Alone

Our earlier group-level results suggest that changes in power (Day 21 – Baseline) provide more information about the state of animal than the amount of power present at Day 21. To test this, we pooled epoch-level data for each metric across animals and labelled each data point as having originated from a non-responder or hyperalgesic animal. The cumulative density functions for changes in power and Day 21 power for each group are shown in Figure 6A and 6B, respectively. These figures clearly show that distributions for non-responders and hyperalgesic animals are more clearly separated when estimates of power change are used. ROC analysis supported this interpretation, with estimates of power change, AUC = .77, *p* < .05, providing more information about class membership than estimates of power amount, AUC = .65, *p* < .05 (Figures 6C & 6D). These AUC values correspond to large (Cohen’s *d* = 1.03) and medium effects (Cohen’s *d* = .51) for power change and power amount, respectively (Rice & Harris, 2005), and the obtained difference between AUC values surpassed the 95^th^ % of a bootstrapped, null distribution (Figure 6E).

**Figure 6.**
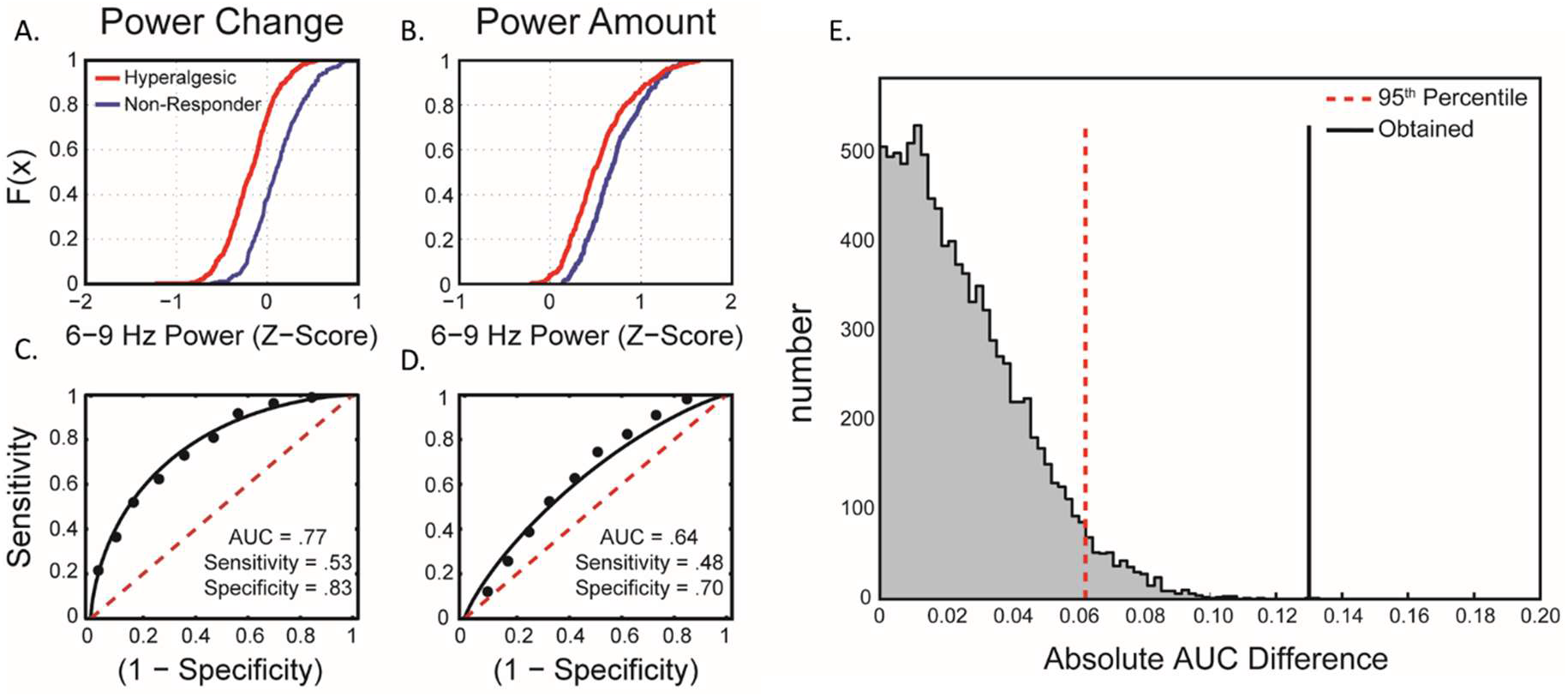
Changes in 6-9 Hz power from Baseline to Day 21 are a better indicator of chronic pain than Day 21 power. **A.** Cumulative density functions of epoch-level 6-9 Hz power changes pooled across animals. For each animal, power changes were calculated by subtracting the average 6-9 Hz power at Baseline from each Day 21 epoch estimate. Hyperalgesic animals are clearly shifted to the left of non-responder animals, indicating that larger decreases in power are associated with hyperalgesia. **B.** Cumulative density function of pooled, epoch-level 6-9 Hz power on Day 21. Hyperalgesic animals are shifted to the left of non-responder animals, but less so than what is seen with power change. ROC analyses of power change (**C**) and Day 21 power (**D**). **E.** The obtained difference in AUC values surpasses the 95^th^% of a null distribution of absolute, AUC differences thereby indicating that power change is significantly more accurate than power amount.

## Discussion

Discovery of biomarkers that can reliably signal the presence of chronic pain, and the risk of transition from acute to chronic pain, are key to improving diagnosis and treatment. Here, we tested whether oscillatory signal changes in response to a preclinical model of neuropathic, orofacial pain could be used as such a biomarker. Following CCI-ION, heterogenous changes to power in the ~6-9 Hz range occurred that were dependent on the behavioral profile of the animal. Specifically, mechanical threshold decreases (i.e. hyperalgesia) on the side of the face ipsilateral to CCI-ION corresponded to *decreases* of 6-9 Hz power, while threshold increases (i.e. non-response) corresponded to *increases* in power. In contrast, we were unable to identify changes to oscillatory peak frequency after injury. Across animals, 6-9 Hz power was significantly correlated with mechanical threshold changes on the afflicted side of the face. Similarly, after accounting for development of contralateral mechanical hyperalgesia in non-responder animals, changes in facial grimace scores—a marker of ongoing pain—were also correlated with spectral power changes in this range. Using ROC analyses in combination with permutation testing, we concluded that power decreases were a reliable marker of chronic pain development in over half of hyperalgesia animals, while power increases were present in the majority of non-responder animals. Finally, we show that power changes provide a more accurate metric than Day 21 power for distinguishing hyperalgesic and non-responder animals.

These findings contrast previous reports of oscillatory changes in the rat brain after injury. Notably, *increases* in Theta range power have been found following either injection of capsaicin into the hindpaw (LeBlanc et al., 2014) or chronic constriction injury of the sciatic nerve (LeBlanc et al., 2016). Similar results have also been described in studies of brief, phasic pain (i.e. Devonshire et al., 2015). One explanation for this discrepancy rests upon differences in how analyses were performed. While we used a modelling approach to disentangle narrowband spectral activity from aperiodic signals, prior studies have not done so. As others have pointed out, failure to account for aperiodic signals can lead to overestimation of oscillatory changes (Haller et al., 2018). However, we found no examples in the current study where the direction of results reversed (i.e. a power decrease became a power increase) when aperiodic modelling was not employed.

Another source of incongruency may relate to definition of the frequency bands of interest. Most studies have relied on traditional, human-derived frequency band definitions, whereas we empirically defined our oscillation of interest. This oscillation encompassed both Alpha and Theta ranges, which some studies have reported respond differently to phasic pain. For example, one recent set of findings indicate that Delta/Low Theta oscillations decrease with noxious stimulation while Upper Theta/Alpha oscillations increase (Peng et al., 2018).

Another possible explanation is that studied oscillations are fundamentally different due to ongoing behaviors that were present when ECoG was recorded. Previous studies have focused on periods of “quiet wakefulness (i.e. LeBlanc et al., 2014) whereas our recordings involved active cage exploration. Similarly, the anatomical origin of injury (orofacial vs. somatic) may be responsible for differences via activation of distinct ascending, nociceptive pathways (trigeminal vs. spinal pathways; Meier et al., 2014; Schmidt et al., 2015; Akintola et al., 2017).

Finally, differences in the sites from which signals were recorded from may play a role. Whereas prior studies have recorded signals from probes placed in either somatosensory cortex (S1) or thalamus (e.g. LeBlanc et al., 2014), we recorded over an area putatively corresponding to the anterior cingulate cortex (ACC). Shih and colleagues (2019) have recently demonstrated that ACC theta power is decreased, while motor cortex theta power is increased following lesion of the medial thalamic nuclei, a region thought to convey affective elements of pain. Likewise, decreases in Theta and Alpha power following application of a noxious, mechanical probe are evident in the ACC but not S1 (Xiao et al., 2019). Thus, discrepancies in the direction of oscillatory changes may reflect real, physiological differences in how distinct brain regions respond to persistent pain. Caution should, however, be taken when attempting to draw firm conclusions based on potential regional differences; ECoG samples neuronal activity at the mesoscopic scale, making it impossible to determine the extent to which recorded oscillatory changes originate from or represent processes in the ACC.

Regardless of the source of these differences, our data demonstrate that changes to oscillations *at the individual animal level* can be used to identify cases of chronic pain. Indeed, we were able to identify meaningful post-CCI changes in more than 60% of animals. To our knowledge, this has not been previously explored in biomarker studies of animal in chronic pain. By comparison, we were unable to identify differences between non-responder and hyperalgesic animals at Day 21 at the group-level data. This suggests that it is difficult to identify aberrant activity from post-injury spectral power alone. This finding was not entirely unexpected, since differences in confounding physiological factors, such as cerebrospinal fluid thickness (Rice et al., 2013), are known to contribute heavily to between-subject variability. From a clinical perspective, these findings suggest that changes in EEG characteristics are likely to be the best diagnostics for detecting chronic pain, and, more perhaps more importantly, for *predicting susceptibility to transition from acute to chronic pain*. Indeed, our ROC analyses indicate that power changes are more sensitive to behavioral outcome than Day 21 power alone. From a clinical perspective, regular collection of EEG measurements during routine physicals would ensure that relevant reference points exist when evaluating future complaints of persistent pain. This might be of particular importance for individuals at greater risk for developing chronic pain, such as fighter-warriors, athletes, and individuals planning elective surgery.

A potential limitation of the current study is the absence of a formal control group of animals. Given our interest in identifying within-animal changes following CCI-ION, we elected to forego this group in order to maximize the number of injured animals included in our study. While we cannot completely rule out the possibility that our results are due to confounds that are correlated with development of chronic pain, such as aging between Baseline and Day 21, we take some comfort in the identified differences between hyperalgesic and non-responder animals. Similarly, analysis of animal activity suggests that spectral power changes cannot be explained simply by increases or decreases in activity across the two timepoints. What these differences reflect is difficult to pinpoint, however, as data from non-responder animals is itself difficult to interpret. Specifically, increases in mechanical thresholds on the injured side may result from a myriad of factors including CCI-ION induced hypoesthesia (i.e. Akintola et al., 2017) or how a given animal responds to repeated testing of injured tissue. Despite this, factors independent of CCI-ION, such as aging, are identical across groups and therefore unlikely to explain the current findings. The strong correlation between spectral and behavioral changes, but not changes to animal activity, provides some comfort that our results reflect a consequence of some element related to persistent pain. Nonetheless, the ultimate test of our findings is whether they can be applied successfully to a novel dataset. To that end, we encourage those interested in using our analytic tools to contact us so that we may share them.

In conclusion, we present novel data demonstrating that decreases in ~6-9 Hz power are not only a reliable marker for detecting chronic pain but also a more sensitive marker than raw spectral power. While future studies are needed to directly address differences between our findings and those described previously, these results provide strong evidence that oscillatory changes hold promise for identifying chronic pain in the clinic.

## Acknowledgements

The authors declare no other conflicts of interest. Research reported in this publication was supported by the National Institute of Neurological Disorders and Stroke of the National Institutes of Health grants R01NS099245 and R01NS104297 (to AK). The content is solely the responsibility of the authors and does not necessarily represent the official views of the National Institutes of Health. The funding sources has no role in study design; the collection, analysis and interpretation of data; the writing of the report; or in the decision to submit the article for publication.

